# Dietary xanthan gum alters antibiotic efficacy against the murine gut microbiota and attenuates *Clostridioides difficile* colonization

**DOI:** 10.1101/786335

**Authors:** Matthew K. Schnizlein, Kimberly C. Vendrov, Summer J. Edwards, Eric C. Martens, Vincent B. Young

## Abstract

Dietary fiber provides a variety of microbiota-mediated benefits ranging from anti-inflammatory metabolites to pathogen colonization resistance. A healthy gut microbiota protects against *Clostridioides difficile* colonization. Manipulation of these microbes through diet may increase colonization resistance to improve clinical outcomes. The primary objective of this study was to identify how the dietary fiber xanthan gum affects the microbiota and *C. difficile* colonization.

We added 5% xanthan gum to the diet of C57Bl/6 mice and examined its effect on the microbiota through 16S rRNA-gene amplicon sequencing and short-chain fatty acid analysis. Following either cefoperazone or an antibiotic cocktail administration, we challenged mice with *C. difficile* and measured colonization by monitoring colony-forming units.

Xanthan gum administration associates with increases in fiber degrading taxa and short-chain fatty acid concentrations. However, by maintaining both the diversity and absolute abundance of the microbiota during antibiotic treatment, the protective effects of xanthan gum administration on the microbiota were more prominent than the enrichment of these fiber degrading taxa. As a result, mice that were on the xanthan gum diet experienced limited to no *C. difficile* colonization.

Xanthan gum administration alters mouse susceptibility to *C. difficile* colonization by maintaining the microbiota during antibiotic treatment. While antibiotic-xanthan gum interactions are not well understood, xanthan gum has previously been used to bind drugs and alter their pharmacokinetics. Thus, xanthan gum may alter the activity of the oral antibiotics used to make the microbiota susceptible. Future research should further characterize how this and other common dietary fibers interact with drugs.

**IMPORTANCE:** A healthy gut bacterial community benefits the host by breaking down dietary nutrients and protecting against pathogens. *Clostridioides difficile* capitalizes on the absence of this community to cause diarrhea and inflammation. Thus, a major clinical goal is to find ways to increase resistance to *C. difficile* colonization by either supplementing with bacteria that promote resistance or a diet to enrich for those already present in the gut. In this study, we describe an interaction between xanthan gum, a human dietary additive, and the microbiota resulting in an altered gut environment that is protective against *C. difficile* colonization.

## INTRODUCTION

The microbiota plays an integral role in gut health by aiding in digestion and regulating colonic physiology (1, 2). Manipulating the microbiota to improve human health by either administering live bacteria (i.e. probiotics) or adding nondigestible, microbiota-accessible ingredients to the host’s diet (i.e. prebiotics) has become a prominent area of biomedical research. While probiotics rely on exogenously added microbes for their effect, diet modification uses indigenous microbes already present in the gut to generate the beneficial effects described above. While the community as a whole may remain intact, diet modification can affect subsets of the community that are better suited to utilize the altered nutrient composition (3). This effect is most prominent in hunter-forager societies where seasonal dietary changes modulate the microbiota (4). In Western diets, a great emphasis has been placed on the types and abundance of host indigestible fiber polysaccharides that are only accessible by the microbiota, such as resistant starch, inulin or the fibers naturally present in fruits, vegetables and whole grains.

Dietary fiber promotes microbial short-chain fatty acid (SCFA) production. While SCFA profiles are unique from individual to individual, they provide a variety of benefits including increased colonic barrier integrity and decreased inflammation (5-10). Depending on the structure of the fiber backbone and side chains, polysaccharides select for unique taxa and as a result, unique fermentation profiles (11). Several key species may be responsible for degrading the fiber’s carbohydrate structure, the byproducts of which go on to be metabolized by a number of additional taxa (12). Butyrate, a short chain fatty acid and product of fiber degradation, has been linked to increased gut barrier integrity and decreased inflammation (13-15). Fiber degradation and SCFA production are also associated with clearance of *Clostridioides difficile*, formerly known as *Clostridium difficile*, following fecal microbiota transfer (FMT) (16). Switching mice to a high fiber diet while colonized with *C. difficile* increased SCFA concentrations and also cleared the infection (17). Since *C. difficile* infection represents a significant healthcare burden, characterizing how these polysaccharides shape the gut environment and impact *C. difficile*’s ability to colonize will provide insight into how they might be used to improve patient outcomes.

Some polysaccharides included in food are added to alter texture rather than for nutritional benefit. Xanthan gum, synthesized by the bacteria *Xanthamonas campestris*, is a common food additive used as a thickener, particularly in gluten free foods, where industrial production is worth approximately $0.4 billion each year. Xanthan gum structure consists of (1→4)-linked *β*-D-glucose with trisaccharide chains containing two mannose and one glucuronic acid residues linked to every other glucose molecule in the backbone, with possible acetylation on the first branching mannose and 3,6-pyruvylation on the terminal mannose (18). These negatively charged side chains give xanthan gum its viscous, gel-like properties. Although not specifically included in foods for their potential prebiotic activity, bacteria can degrade xanthan gum to increase fecal SCFA concentrations (19, 20). However, little is known about what bacterial taxa are involved in these transformations.

This study investigated the effect of xanthan gum on the bacterial composition of specific pathogen-free C57Bl/6 mice and its effect during an antibiotic model of *C. difficile* infection. Our goal for this paper was to (i) characterize the effects of xanthan gum on the mouse microbiota and (ii) characterize the effects of xanthan gum on *C. difficile* colonization. Surprisingly, we found that xanthan gum administration alters mouse susceptibility to *C. difficile* colonization by maintaining the microbiota during antibiotic treatment.

## RESULTS

### Xanthan gum maintains the abundance of microbial taxa during cefoperazone treatment

Using C57Bl/6 mice, we tested the effects of xanthan gum on the microbiota using mouse models designed to study the effects of antibiotic perturbation. Since our initial goal was to study the effects of xanthan gum on *C. difficile* infection in mice, these models entailed multiple days of antibiotic treatment necessary to make the microbiota susceptible to *C. difficile* (Fig. 1A, Fig. S1A). Some mice were kept on a standard mouse chow diet; the rest were put on an equivalent diet supplemented with 5% xanthan gum.

**Figure 1.**
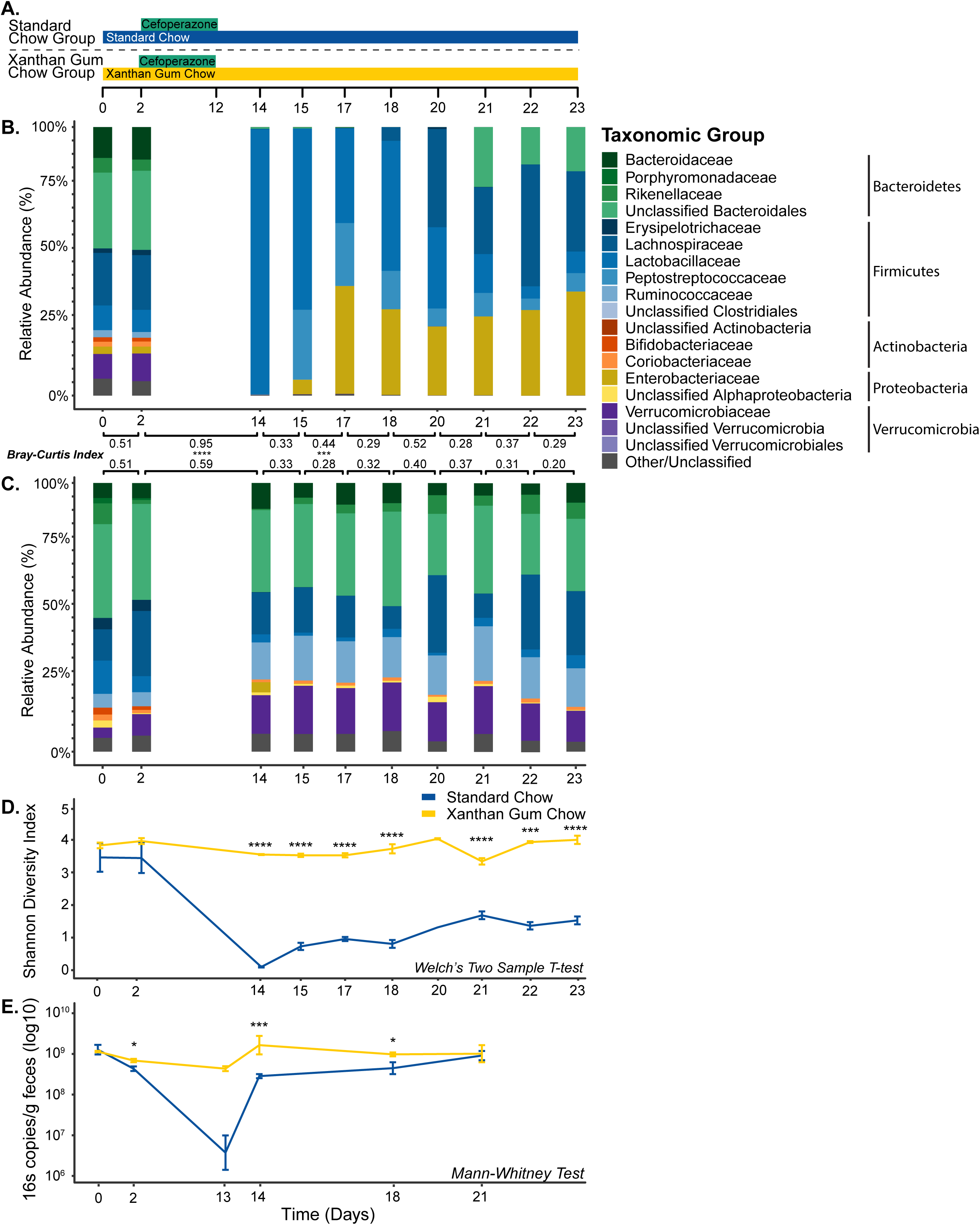
Fecal bacterial diversity and abundance during xanthan gum and cefoperazone administration. **A**. Timecourse of the experimental model for the mice on standard and xanthan gum chows. **B**. Microbiota mean relative abundance in mice on standard chow (N = 5). **C**. Microbiota mean relative abundance in mice on xanthan gum chow (N=6). Bray-Curtis dissimilarity index is shown comparing each timepoint. **D**. Mean Shannon diversity index of the bacterial communities shown in B. and C. (error bars indicate 1 std dev.). Statistical testing with Welch’s Two-Sample T-test. **E**. Bacterial absolute abundance indicated by qPCR using “universal” primers for the 16S rRNA-gene (normalized to g of feces; error bars indicate 1 std. dev.). Statistical analysis: Mann-Whitney for β-diversity and 16S qPCR as well as Welch’s Two-Sample T-test for Shannon Diversity (* indicates p < 0.05; ** p < 0.01, *** p < 0.001, **** p < 0.0001).

In the cefoperazone mouse model, 16S rRNA-gene analysis of mouse fecal samples revealed a baseline microbiota dominated by Bacteroidetes (∼45%) and Firmicutes (∼35%), with the remainder of the community composed of Actinobacteria, Proteobacteria and Verrucomicrobia (Fig. 1B & C). Following cefoperazone treatment of mice on standard chow, Lactobacillaceae predominated a fluctuating community, as evidenced by increased mean Bray-Curtis distances between timepoints. While we also observed higher dissimilarity in the xanthan gum chow group following antibiotics, microbial communities were significantly more similar in the xanthan gum chow group compared to the standard chow group, as measured by Bray-Curtis distances. However, the relative abundance of bacterial taxa remained similar after cefoperazone treatment in mice fed 5% xanthan gum (Fig. 1C). These protective effects are reflected in a significantly higher Shannon Diversity and absolute abundance of fecal bacteria in xanthan gum-fed mice following cefoperazone treatment compared to those on standard chow (Fig. 1D & E). These data suggest that high concentrations of xanthan gum prevent cefoperazone-mediated alterations to the mouse microbiota. To see if xanthan gum had a similar protective effect for other antibiotics, we also used an oral antibiotic cocktail model coupled with intraperitoneal clindamycin, which has also been shown to render mice susceptible to *C. difficile* colonization (21, 22). However, the microbiota differences between chow groups were less pronounced (Fig. S1B & C). Taken together, our results show that xanthan gum administration maintains both diversity and overall abundance of microbes in the gut during cefoperazone treatment.

Using the linear discriminant analysis algorithm LEfSe, we identified 35 OTUs that were significantly increased two days following the switch from standard to xanthan gum chow (Fig. S3). We also observed a shift in bacterial metabolism marked by significantly higher butyrate and propionate concentrations in mice on xanthan gum chow compared to those on standard chow (Fig. S4). No significant OTUs were identified comparing the same timepoints in the standard chow group. Following cefoperazone treatment, 4 OTUs were increased and 80 OTUs were decreased in the xanthan gum group (Fig. S5). In the standard chow group, only 1 OTU (Lactobacillus) significantly increased following cefoperazone treatment (Fig. S6). Unsurprisingly, 48 of the 112 OTUs that were negatively correlated with cefoperazone treatment in the standard chow group were also negatively correlated in the xanthan gum group.

### Xanthan gum-mediated microbiota protection limits *C. difficile* colonization

Two days after the mice were removed from cefoperazone, they were challenged with *C. difficile* strain 630g spores administered by oral gavage. By monitoring feces for colony-forming units (both vegetative cells and spores), we observed approximately 1×10^6^ CFU/g feces *C. difficile* in mice on standard chow 1 day post-gavage (Day 15), which rose to 1×10^7^ for the duration of the experiment (Fig. 2). However, when on xanthan gum, only a small number of CFU was observed 1 day post-gavage but by day 4 (Day 19) all mice had cleared *C. difficile* (Fig. 2). In the antibiotic cocktail model, *C. difficile* colonized mice on both standard and xanthan gum chow similarly until 7 days post-gavage (Day 15) when *C. difficile* colonization levels were significantly lower in the mice on xanthan gum chow (Fig. S7).

**Figure 2.**
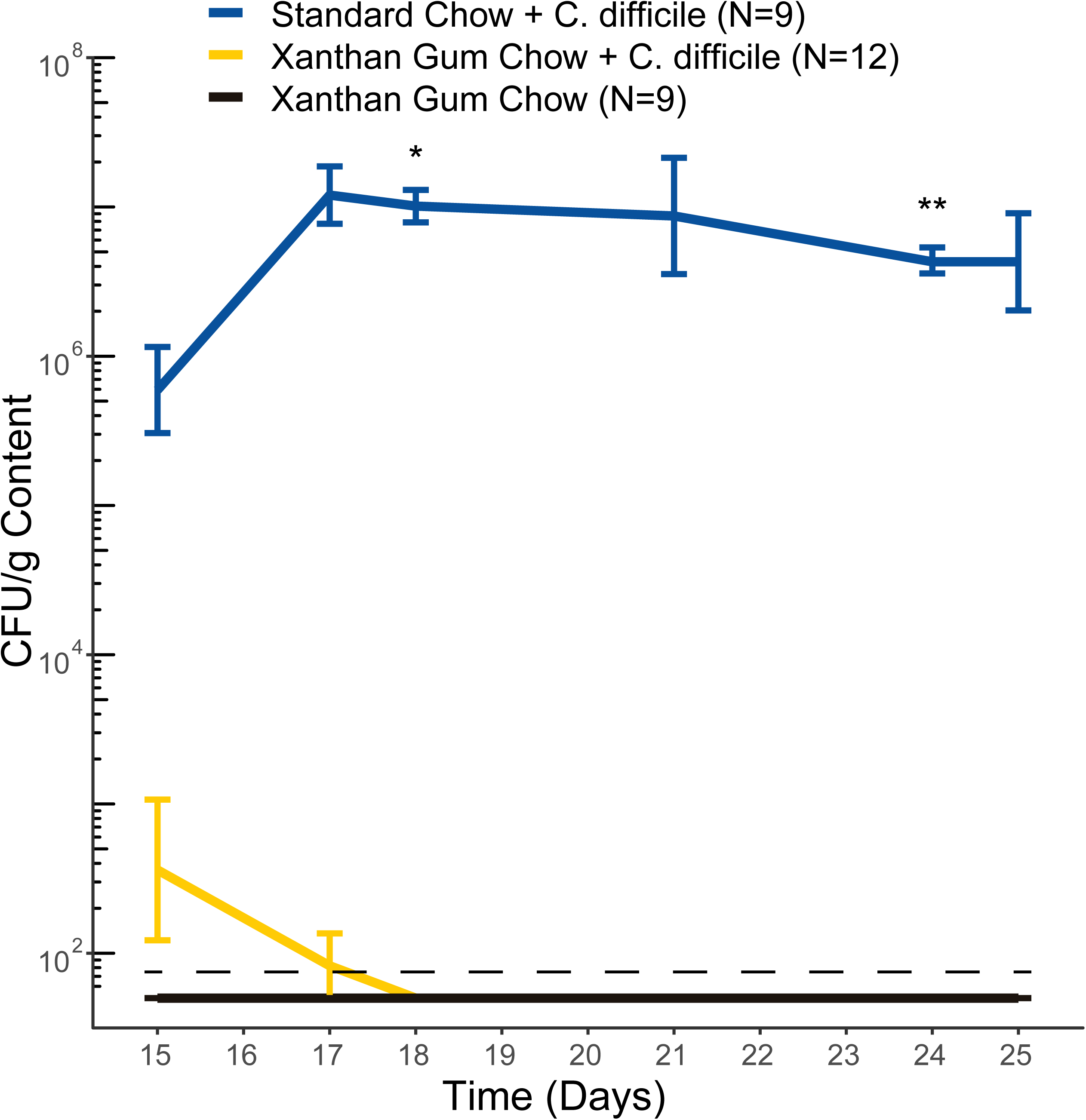
*C. difficile* colonization in mice on standard and xanthan gum chows. *C. difficile* colony-forming units (CFUs) in cefoperazone treated mice were normalized to fecal mass. Lines indicate mean CFU levels (error bars indicate 1 std dev). Data shown are from both experiments 1 and 2. Statistical testing was performed using Welch’s 2 Sample T-Test (* indicates p < 0.05; ** p < 0.01).

## DISCUSSION

The use of dietary polysaccharides for their beneficial health effects, either directly on the host or indirectly through the microbiota, has been widely demonstrated (15, 19). In the context of *C. difficile*, diet may play a role in pathogen evolution, such as with trehalose, or influence colonization resistance, such as with dietary fiber and zinc (17, 23-25). Dietary alteration may shape the intestinal environment by altering the nutrients available or by modulating the concentrations of compounds toxic to *C. difficile*, such as secondary bile salts. As a common food additive, xanthan gum’s physicochemical properties are well known (18). However, its impacts on the gut microbiota are poorly understood. Although we were not able to test whether xanthan gum enriches for fiber-degrading bacteria to increase colonization resistance, we did observe that xanthan gum interferes with the activity of orally administered antibiotics to protect mice from *C. difficile* colonization. These protective effects vary by type of antibiotic. While xanthan gum may have enriched for taxa capable of degrading it, these changes were minor compared to the much larger differences observed between diet groups during antibiotic treatment.

As a third-generation cephalosporin, cefoperazone has broad-spectrum efficacy (26, 27). As a result, it is not surprising that, in the standard chow group, it had a significant impact on microbiota community structure. These results agree with previously published work on the effect of cefoperazone on the murine gut microbiota (28, 29). As demonstrated in this study, diet can affect antibiotic efficacy in unexpected ways. While both bacterial diversity and abundance were maintained in mice on xanthan gum, the similarities in OTUs identified by LEfSe between the two groups indicates that cefoperazone affected the microbiota in both groups but was attenuated in the xanthan gum chow group. Since we observed similar antimicrobial activity against an *Escherichia coli* strain ECOR2 lawn from fecal extracts obtained during antibiotic administration between diet groups and no inhibitory activity in fecal extracts from non-antibiotic treated control mice (data not shown), our data suggests that while still active, cefoperazone’s efficacy was interrupted *in vivo*.

By at least partially protecting the microbiota from the effects of cefoperazone, xanthan gum administration preserved colonization resistance to *C. difficile*. Colonization resistance comprises a variety of mechanisms including the metabolism of bile salts and competition for nutrients (30). Microbially-modified secondary bile salts inhibit *C. difficile* outgrowth much more than their primary precursors (31). Microbial metabolism mediates a variety of modifications to primary bile salts, including deconjugation by *Lactobacillus* and *Bifidobacterium* sp. as well as 7α-dehydroxylation by *Clostridium* sp. (32-35). The lack of secondary metabolites produced by these taxa has been correlated with a lack of colonization resistance (31, 36-38). The indigenous microbiota also prevents *C. difficile* from establishing itself within the colonic environment by limiting the nutrients available for growth (39, 40). A number of taxa, including the Lachnospiraceae, have been shown to provide resistance to *C. difficile* colonization, which may occur through niche competition (41, 42). Despite increased SCFA concentrations immediately following xanthan gum administration, direct alterations of the microbiota by xanthan gum did not appear to affect colonization resistance on the day of *C. difficile* gavage since SCFA concentrations had returned to baseline levels. By protecting the microbiota during antibiotic treatment, xanthan gum likely maintained these metabolic mechanisms to exclude *C. difficile* from the gut. This suggested that while the community was altered, enough bacterial taxa remained to exclude *C. difficile*.

While we did not demonstrate a mechanism for xanthan gum’s effect, its gel-like nature may interrupt the activity of antibiotics by altering their pharmacokinetics. Several large polysaccharides with negatively charged or polar sidechains, such as hydroxypropylmethyl cellulose, mannan oligosaccharides and guar gum, increase the excretion of cholesterol and bile salts in feces by limiting their absorption (43-49). While not previously reported, xanthan gum may also interact with these compounds. Similarities between the chemical structures of these sterol ring-containing compounds and of cefoperazone may result in interactions between xanthan gum and the antibiotic. The greater efficacy of the antibiotic cocktail plus clindamycin model against the microbiota is likely due to varied interactions with the five antibiotics in addition to the effect of the intraperitoneal injection of clindamycin. While potential alterations to the bile salt pool by xanthan gum may have limited *C. difficile* germination, we observed more fecal CFUs 1 day post-gavage (day 15) than what we used to inoculate the mice on day 14, suggesting that any disruptions to enterohepatic circulation did not prevent germination as there was some vegetative cell outgrowth. Furthermore, we have previously observed that few spores (i.e. <100) are sufficient to infect mice, suggesting that it was not a small inoculum that protected the mice (unpublished data).

Polysaccharide-drug interactions are frequently explored as means to delay drug release *in vivo*. When mice consume xanthan gum in their chow, orally administered antibiotics may become trapped inside the gel formed by hydrated xanthan gum. Previous research has shown that xanthan gum would provide time-dependent release that occurs at a slower rate than other large, polar polysaccharides. For example, hydroxypropylmethyl cellulose requires three times the concentration to achieve similar drug binding levels as xanthan gum (50, 51). The binding affinity of xanthan gum is pH-dependent, where higher pH limits drug release due to increased integrity of the polymer structure (51). Furthermore, environments with higher ionic strength as well as the presence of other polysaccharides increases xanthan gum’s drug retaining efficiency (52, 53). Thus, the colonic environment would be conducive for high xanthan gum affinity for binding compounds such as cefoperazone due to its relatively higher pH.

In our study, dietary xanthan gum administration protected the microbiota during antibiotic treatment leading to the exclusion of *C. difficile* from the gut. While our study suggests that a common dietary polysaccharide interacts with the effects of antibiotics, there are several limitations that merit future research. Since few individuals will consume xanthan gum at the concentrations we used, titering in lower doses of xanthan gum to get closer to physiological levels would elucidate the effects of xanthan gum in a normal human diet. Future research should also characterize how polar polysaccharides such as xanthan gum interact with compounds in the gut. This would be important for understanding drug pharmacokinetics as well as the impact of xanthan gum on bile salts and enterohepatic circulation. Further work characterizing this common food additive would provide a greater understanding not only of how it is degraded in the gut it but also the potential positive effects of its fermentative byproducts.

## METHODS

### Ethics statement

The University Committee on Use and Care of Animals of the University of Michigan, Ann Arbor, approved all animal protocols used in the present study (PRO00008114). These guidelines comply with those set by the Public Health Service policy on Humane Care and Use of Laboratory Animals.

### Animals and housing

We obtained five-to eight-week old, male and female mice from an established breeding colony at the University of Michigan. These mice were originally sourced from Jackson Labs. We housed mice in specific pathogen-free and biohazard AALAC-accredited facilities maintained with 12 h light/dark cycles at an ambient temperature of 22 ±2°C. All bedding and water were autoclaved. Mice received gamma-irradiated food (LabSupply 5L0D PicoLab Rodent Diet, a gamma-irradiated version of LabSupply 5001 Rodent LabDiet) or an equivalent diet with 5% xanthan gum added (95% LabSupply 5001 Rodent LabDiet, 5% xanthan gum [Sigma]; gamma-irradiated by manufacturer). We housed mice in groups of two to five animals per cage, with multiple cages per treatment group.

All cage changes, infection procedures, and sample collections were conducted in a biological safety cabinet (BSC) using appropriate sterile personal protective equipment between cage contacts. The BSC was sterilized with Perisept (Triple S, Billerica, MA) between treatment groups. Gloves were thoroughly sprayed with Perisept between each cage and completely changed between groups. A description of the metadata for the mouse experiments including cage number and treatment group can be found in Table S1.

### Xanthan gum-cefoperazone mouse model

To investigate the effect of xanthan gum on cefoperazone treated mice, we switched mice to a diet containing 5% xanthan gum on day zero. Two days later, we gave mice 0.5 mg/mL cefoperazone in the drinking water for 10 days as previously described to render the mice susceptible to *C. difficile* colonization (54, 55). We changed the antibiotic-water preparation every 2 days. Following 10 days of cefoperazone, we switched mice to Gibco distilled water. We orally gavaged mice with between 10^2^ and 10^4^ *C. difficile* 630g spores or vehicle control (sterile water) 2 days after removing the mice from antibiotics. Spores were prepared as previously described and then suspended in 200 uL of Gibco distilled water and heat-shocked (55). Viable spores were quantified immediately after gavage using taurocholate cycloserine cefoxitin fructose agar (TCCFA) as previously described (55).

### Xanthan gum-antibiotic cocktail mouse model

To investigate the effect of xanthan gum on an alternative antibiotic model (antibiotic cocktail with clindamycin), we switched mice to a 5% xanthan gum diet on day zero and then put on an antibiotic cocktail (0.4 mg/mL kanamycin, 0.035 mg/mL gentamicin, 850 U/mL colistin, 0.215 mg/mL metronidazole, and 0.045 mg/mL vancomycin) for 3 days in their drinking water as previously described (21, 22). On day 5, we removed mice from oral antibiotic administration and returned them to regular drinking water. On day 7, mice were given an intraperitoneal (IP) injection of clindamycin hydrochloride (10 mg/kg). 1 day following the ip injection, we orally gavaged mice with between 10^2^ and 10^4^ *C. difficile* 630g spores as described above.

### Quantitative culture

We suspended fresh fecal pellets in sterile, pre-reduced Gibco PBS (ThermoFisher) using a ratio of 1 part feces to 9 parts Gibco PBS, wt/vol (ThermoFisher, Waltham, MA). We serially diluted these suspensions, plated them on TCCFA, and incubated the plates anaerobically at 37°C for 18-24 hrs before counting colonies.

### 16S rRNA-gene Qpcr

We suspended fecal pellets in PBS as described above and centrifuged them at 6,000 rpm for 1 minute. 100-400 uL of supernatant was removed for metabolite analysis. Using the sedimented fecal content, we performed DNA extractions using the DNeasy PowerSoil Kit (Qiagen, Germantown, MD), according to the manufacturer’s protocol. We immediately stored extracted DNA at −20°C until further use. We then performed qPCR on a LightCycler® 96 thermocylcer (Roche, Basel, Switzerland) using the PrimeTime® Gene Expression Master Mix (IDT, Coralville, IA) and a set of broad range 16S rRNA gene primers (56). All fecal DNA was amplified in triplicate with *Escherichia coli* genomic DNA standards in duplicate and negative controls in triplicate. The LightCycler reaction conditions were as follows: 95°C for 3 minutes, followed by 45 cycles of 2-step amplification at 60°C for 60 s and 95°C for 15 s. C_q_ values for each reaction were determined using the LightCycler® 96 software, and fecal DNA concentrations were determined by comparing C_q_ values to the standards in each plate and normalizing to each individual sample’s fecal mass. We used Welch’s 2-Sample T-Test to test for significance.

### Short-chain fatty acid analysis

100 uL of fecal supernatants were filtered at 4°C using 0.22 micron 96-well filter plates and stored at −80°C until analysis. We transferred the filtrate to 1.5 mL screw cap vials with 100 uL inserts for high performance liquid chromatography analysis (HPLC) and then randomized them. We quantified acetate, propionate, and butyrate concentrations using a refractive index detector as part of a Shimadzu HPLC system (Shimadzu Scientific Instruments, Columbia, MD) as previously described (8). Briefly, we used a 0.01 N H_2_SO_4_ mobile phase through an Aminex HPX87H column (Bio-Rad Laboratories, Hercules, CA). Sample areas under the curve were compared to volatile fatty acid standards with concentrations of 40, 20, 10, 5, 2.5, 1, 0.5, 0.25, and 0.1 mM. Through blinded curation, we assessed baseline and peak quality and excluded poor quality data if necessary.

### DNA Extraction and Illumina MiSeq sequencing

The detailed protocol for DNA extraction and Illumina MiSeq sequencing was followed as described in previous publications with modifications (55). Briefly, 200-300 uL of 10x diluted fecal pellet was submitted for DNA isolation using the MagAttract PowerMicrobiome DNA isolation kit (Qiagen, Germantown, MD). Samples were randomized into each extraction plate. To amplify the DNA, we used barcoded dual-index primers specific to the V4 region of the 16S rRNA-gene (57). Negative and positive controls were run in each sequencing plate. Libraries were prepared and sequenced using the 500-cycle MiSeq V2 reagent kit (Illumina, San Diego, CA). Raw FASTQ files, including the appropriate controls, were deposited in the Sequence Read Archive (SRA) database (Accession #’s: XXXXXXXXXX; *Note: the authors have submitted the sequences, but they are still being processed by the SRA*).

### Data processing and microbiota analysis

16S rRNA-gene sequencing was performed as previously described using the V4 variable region and analyzed using mothur. Detailed methods, processed read data, and data analysis code are described on GitHub (https://github.com/mschnizlein/xg_microbiota). Briefly, after assembly and quality control, such as filtering and trimming, we aligned contigs to the Silva v.128 16S rRNA database. We removed chimeras using UCHIME and excluded samples with less than 5000 sequences. We binned contigs by 97% percent similarity (OTU) using Opticlust and then used the Silva rRNA sequence database to classify those sequences. Alpha and beta diversity metrics were calculated from unfiltered OTU samples. We used LEfSe to identify OTUs that significantly associated with changes across diets and antibiotic treatments (58). We performed all statistical analyses in R (v.3.5.2).

## Availability of data

Raw FASTQ files are available on the SRA. Code and detailed processing information, as well as raw data are available on GitHub (https://github.com/mschnizlein/xg_microbiota).

## Acknowledgements

The authors would like to thank the Host Microbiome Initiative and Microbial Systems Molecular Biology Laboratory at the University of Michigan for their support with the 16S rRNA sequencing. We would also like to thank Thomas Schmidt and Kwi Kim for their help with the HPLC analyses and Matt Ostrowski for guidance with the project’s direction. This research was funded by the NIH (5-U01-AI124255).

## Availability of data and analysis material

SRA: Sequences have been deposited. However, the authors are waiting to hear back from the SRA for their bioproject and biosample numbers.

GitHub: https://github.com/mschnizlein/xg_microbiota

## Author’s contributions

MKS, KCV, ECM, VBY – Conception or design of the work

MKS, KCV, SJE – Data collection

MKS, SJE – Data analysis

MKS, SJE – Drafting the article

MKS, KCV, SJE, ECM, VBY – Critical revision of the article

MKS, KCV, SJE, ECM, VBY – Final approval of the version to be published

**Figure S1 Fecal (Days 0-15) and cecal (Day 22) bacterial diversity and relative abundance during xanthan gum and antibiotic cocktail administration. A**. Time course of the experimental model for the mice on standard and xanthan gum chows. **B**. Microbiota mean relative abundance in mice on standard chow (N = 5). **C**. Microbiota mean relative abundance in mice on xanthan gum chow (N=7). Bray-Curtis dissimilarity index is shown comparing each timepoint. **D**. Mean Shannon diversity index of the bacterial communities shown in B. and C. (error bars indicate 1 std dev.). Statistical analyses: Mann-Whitney for β-diversity as well as Welch’s Two-Sample T-test for Shannon Diversity (* indicates p < 0.05; ** p < 0.01, *** p < 0.001, **** p < 0.0001).

**Figure S2 LEfSe analysis of the microbiota of mice before and after the start of xanthan gum administration**. Bacterial taxa that were **A**. increased or **B**. decreased following the switch to xanthan gum. The numbers in parentheses indicate the number of OTUs that fall under that particular taxonomic classification.

**Figure. S3 Short chain fatty acid analysis** (acetate, propionate, and butyrate) of mouse fecal content from the cefoperazone model. Time points show before (Day 0) and after diet change (Day 2) as well as after antibiotic treatment (Days 13 and 14). Statistics were performed using Welch’s Two-Sample T-test. (** indicates p < 0.01).

**Figure S4 LEfSe analysis of the microbiota in mice on xanthan gum chow during cefoperazone treatment**. Bacterial taxa that were **A**. increased **B**. or decreased following cefoperazone treatment. The numbers in parentheses indicate the number of OTUs that fall under that particular taxonomic classification.

**Figure S5 LEfSe analysis of the microbiota of mice on standard chow during cefoperazone treatment**. Bacterial taxa that were **A**. increased **B**. or decreased following cefoperazone treatment. The numbers in parentheses indicate the number of OTUs that fall under that particular taxonomic classification.

**Figure S6 C. difficile CFU in mice in the antibiotic cocktail model**. *C. difficile* colony-forming units (CFUs) in antibiotic cocktail treated mice were normalized to fecal mass. Lines indicate mean CFU levels (error bars indicate 1 std dev). Data shown are from both experiments 3 and 4. Statistical testing was performed using Welch’s 2 Sample T-Test (* indicates p < 0.05).

**Table S1: Mouse experiment and analysis metadata**

**Table S2: Mouse sequencing metadata from the cefoperazone model**

**Table S3: Mouse sequencing metadata from the antibiotic cocktail with clindamycin model**

**Table S4: LEfSe metadata from the comparison of bacterial communities before and after the start of xanthan gum administration (xgdiet) as well as cefoperazone treatment in the xanthan gum (xgcef) and standard chow groups (stdcef)**.

